# Somatosensory gating dysfunction is masked by cognitive variability in cognitively impaired individuals

**DOI:** 10.64898/2026.06.29.735283

**Authors:** Mahak Virlley, Yin Xi, Natalie M. Bell, Tyrell Pruitt, Lin Guo, Sloan White, Fang F. Yu, Laura H. Lacritz, Heidi Rossetti, C. Munro Cullum, Amil M. Shah, Elizabeth M. Davenport, Joseph A. Maldjian, Amy L. Proskovec

## Abstract

Disruptions in somatosensory processing have been observed in cognitive impairment (CI), suggesting that alterations in sensory processing may emerge earlier during cognitive decline than previously recognized. Somatosensory gating (SG) is an automatic inhibitory mechanism that protects neural resources by suppressing responses to redundant, non-behaviorally relevant stimuli. Prior work has demonstrated exaggerated gamma SG and response amplitudes in the primary somatosensory cortex (S1) of individuals with Alzheimer’s disease-confirmed pathology, and these effects were masked by variability in attention/executive function performance. However, whether similar relationships are present during earlier stages of cognitive decline, such as CI, remains unclear. Herein, 63 cognitively healthy older adults (CH; mean age = 59.9 ± 8.6 years) and 32 individuals with CI (mean age = 62.4 ± 8.8 years) underwent magnetoencephalography (MEG) while completing a paired-pulse SG paradigm designed to probe inhibitory sensory processing. MEG oscillatory responses were source-imaged using a beamformer. Time series data were extracted from the peak voxel to quantify oscillatory dynamics and SG. Neuropsychological testing was conducted to assess attention/executive function. After controlling for attention/executive function variance, exaggerated gamma SG was observed in adults with CI compared with CH adults (*p* < 0.05). Additionally, adults with CI exhibited increased beta peak frequency following the second stimulation (*p* < 0.01) and a group-by-age interaction for theta SG in S1 (*p* < 0.05). Together, these results suggest somatosensory abnormalities are present in earlier stages of cognitive decline and highlight a dynamic interaction between sensory processing and cognitive systems during this decline.

## Introduction

Cognitive change is a normal process of aging and encapsulates a complex process in which certain cognitive abilities decline gradually over time, while others are more resilient to or even improve with aging (Arif et al., 2020; Committee on the Public Health Dimensions of Cognitive Aging; Board on Health Sciences Policy; Institute of Medicine; Blazer DG, 2015; Deary et al., 2009; Harada et al., 2013; Murman, 2015; Peters, 2006; Proskovec et al., 2016; Wilson et al., 2016). However, some individuals experience cognitive declines that are more severe than expected for their age (i.e., accelerated or pathological cognitive aging) yet do not significantly disrupt daily functioning, representing a transitional zone between normal aging and dementia (Anand S, 2025). This range of subtle-but-meaningful cognitive decline is clinically captured by the construct of mild cognitive impairment (MCI). MCI can progress to dementia [e.g., Alzheimer’s disease (AD)] at which point there is substantial interference with daily functioning and independence. Importantly, such early cognitive impairment (CI) is heterogeneous across individuals and investigations in a heterogeneous and population-based sample may offer generalizable markers of dysfunction. Specifically, characterizing early neural changes associated with CI is critically important for three primary reasons. First, although pathological changes may already be underway, cognitive and functional decline at this stage may still be amenable to early intervention (Anand S, 2025). Second, quantifiable biomarkers of brain dysfunction provide objective targets for evaluating therapeutic agents (e.g., pharmacological, behavioral, neuromodulation) designed to address functional abnormalities associated with disease processes (Collie & Maruff, 2000; Ewers et al., 2011; Gkintoni et al., 2025; Levy et al., 2022). Third, such markers may guide the development of novel therapies to impede cognitive decline or even regain function.

A growing body of work has demonstrated that sensory processing deficits are prevalent in individuals with CI, with much of the literature focusing on the auditory and visual domains (Anstey et al., 2001; Karrasch et al., 2006; Rhodus et al., 2022; Roberts & Allen, 2016; Zuppichini et al., 2025). Because sensory systems rely on coordinated activity across distributed neural circuits, alterations in sensory processing may reflect broader changes in neural function that accompany cognitive decline. Consistent with this view, sensory deficits have been associated with both increased risk for CI and greater cognitive decline (e.g., dementia) (Baltes & Lindenberger, 1997; Cai et al., 2023; Dintica et al., 2023; Moller et al., 2024; Roberts & Allen, 2016; Schubert et al., 2017). Theoretical frameworks, such as the information degradation hypothesis, propose that impaired sensory processing increases cognitive demands on higher-order systems, including attention and executive function (Monge & Madden, 2016; Roberts & Allen, 2016). As these domains compensate for degraded sensory input, cumulative cognitive load increases, potentially accelerating cognitive decline (Roberts & Allen, 2016). Contrary to earlier assumptions that somatosensory dysfunction emerges only in later stages of dementia, somatosensory processing has been shown to interact with cognitive domains across the lifespan, and tactile sensations have been found to have predictive power in MCI diagnosis (Humes et al., 2013; Ibarra-Castaneda et al., 2025; Loffler et al., 2024). These findings suggest that alterations in somatosensory processing may reflect early changes in neural systems that support both sensory and cognitive function, rather than simply representing downstream consequences of advanced neurodegeneration. Notably, Stephen et al., reported significantly increased somatosensory response amplitudes in the primary somatosensory cortex (S1) in individuals with amnestic MCI compared with healthy controls (Stephen et al., 2010). Moreover, these amplified responses were associated with poorer neuropsychological performance, particularly in attention and executive functioning, suggesting a link between aberrant somatosensory processing and cognitive dysfunction (Stephen et al., 2010). Studying somatosensory inhibitory mechanisms therefore provides a tractable approach for probing early neural dysfunction within sensory systems that interact with higher-order neural processes.

One mechanism through which cognition may influence and mask somatosensory processing deficits in CI is through somatosensory gating (SG), a functional sensory inhibition process assessed using paired-pulse somatosensory stimulation (Cromwell et al., 2008). SG refers to the reduced neural response to a second, identical stimulus (Stim2) delivered in short succession (∼500 ms) relative to an initial stimulus (Stim1), reflecting the brain’s ability to suppress redundant behaviorally-irrelevant sensory information (Cromwell et al., 2008). SG primarily occurs in S1 (Cheng et al., 2016; Onishi et al., 2018; Spooner et al., 2019; Wiesman et al., 2017; Wiesman & Wilson, 2020). Functionally, SG is thought to preserve limited neural resources by filtering repetitive sensory input, thereby preventing unnecessary engagement of higher-order cognitive systems (Cromwell et al., 2008). Consistent with this role, disruptions in sensory gating have been associated with adverse sensorimotor and cognitive outcomes, highlighting its importance for both neural efficiency and behavior (Arpin et al., 2017; Heinrichs-Graham et al., 2023; Liu et al., 2018; Pesonen et al., 2023; Spooner et al., 2021; Wiesman et al., 2021; Wiesman & Wilson, 2020).

In contralateral S1, SG of gamma (30+ Hz) neural oscillatory responses has been shown to decline with healthy cognitive aging (Proskovec et al., 2020; Spooner et al., 2019). Interestingly, a prior study reported aberrant gamma SG in contralateral S1 of patients along the Alzheimer’s spectrum (i.e., a combined group of individuals with amnestic MCI or Alzheimer’s dementia) (Wiesman et al., 2021). Specifically, gamma SG was found to be exaggerated in these patients, indicating increased inhibitory mechanisms in S1, but only when performance on attention/executive function and processing speed measures were accounted for (Wiesman et al., 2021). That is, variability in attention/executive function and processing speed masked somatosensory dysfunction along the Alzheimer’s spectrum. Furthermore, in healthy controls, alpha, beta, and theta SG are modulated during engagement of attentional processes (Wiesman & Wilson, 2020). Alpha SG was also aberrant in individuals along the Alzheimer’s spectrum, but variability in cognitive performance did not impact this effect (Wiesman et al., 2021). Finally, SG of evoked neural responses did not differ between individuals along the Alzheimer’s spectrum and cognitively healthy adults, regardless of whether cognitive variability was considered, emphasizing the importance of evaluating induced (i.e. oscillatory) SG (David et al., 2006; Wiesman et al., 2021). Overall, the study conducted by Wiesman et al. emphasizes the importance of considering between-participant variability in neurocognitive decline when probing for dysfunctional sensory processing in dementia (Wiesman et al., 2021). However, it remains unclear whether alterations in gamma SG are already present during the earlier stages (i.e., CI), prior to progression to dementia. Clarifying this relationship may improve our understanding of how early neural changes in sensory inhibition relate to cognitive decline.

In the present study, we extend prior work on somatosensory processing in cognitive decline by examining SG in individuals with CI relative to cognitively healthy (CH) controls. Building on the findings from dementia populations (Wiesman et al., 2021), we evaluated whether a similar masking effect was present in individuals with CI by testing differences in gamma SG and response amplitude both before and after accounting for attention and executive functioning or processing speed, relative to CH controls. For completeness, we also conducted parallel analyses in the theta, alpha, and beta frequency bands. In addition, several exploratory analyses were conducted. First, because prior studies have reported shifts in peak frequency of neural oscillations in CI (Garces et al., 2013), we examined whether the peak frequency of each oscillatory response differed between individuals with CI and CH controls. Second, given evidence of abnormal spontaneous activity in CI (Jiang et al., 2025), we assessed whether spontaneous activity within each frequency band differed by group. Finally, prior work has shown that age interacts with clinical status to influence SG outcomes in CI populations (Spooner et al., 2020). Accordingly, we examined whether age-related changes in SG differed between individuals with CI and CH controls.

## Methods

### Study and ethics

This study utilized data from the Dallas Hearts and Minds Study (DHMS) and the Healthy and Pathological Neurocognitive Aging (HPNA) study. The DHMS is a large, longitudinal, population-based investigation designed to examine cardiovascular health and, more recently, neurocognitive function, with intentional racial minority enrichment. Each study was reviewed and approved by the Institutional Review Board at the University of Texas Southwestern Medical Center, and all participants provided written informed consent following a thorough description of study procedures. Within DHMS, MEG data were acquired in only a subset of participants due to standard contraindications, including the presence of ferromagnetic implants, and scheduling constraints on the machine (e.g., prioritization of clinical epilepsy scans in the mornings). To address this limitation and increase the number of participants from the DHMS cohort with MEG somatosensory gating (SG) data, the HPNA study was conducted as an extension of DHMS. HPNA followed the identical MEG and brain MRI protocols (see DHMS MEG protocol (Bell et al., 2026)), enabling harmonization of data across studies, and focused on the recruitment of individuals from clinical populations of interest (e.g., CI). Across studies, data collection also included demographic information, medical history, and neurocognitive assessments. Exclusion criteria for the present analysis included neurological (beyond CI) or current neuropsychiatric disorders, recent concussion (within the past year), current high-intensity alcohol use (Hingson et al., 2017), or any medical condition affecting the central nervous system.

### Group definitions and participants

A total of 319 DHMS participants completed the MEG SG paradigm, of whom 217 were classified as non-impaired, 92 were classified as impaired, and seven deferred (e.g., non-English speaker). Cognitive status of each of the participants was determined by consensus amongst two board-certified neuropsychologists who reviewed demographic, medical history, and neurocognitive performance data (see *Neuropsychological Testing* below for further details).

The impaired group was further stratified into individuals with dementia (N = 3), individuals with CI likely due to factors other than neurodegenerative disease (e.g., schizophrenia, severe depression; CI-Other; N = 33), and those with CI not confounded by other significant disease processes (N = 59). Individuals with dementia or CI-Other (CI due to other medical comorbidities) were excluded from the CI group analyses. From the CI group, additional participants were excluded due to medical comorbidities (n = 23), incomplete demographic data (n = 4), or excessive MEG artifact (n = 2). To increase the CI group sample size, two additional participants from the HPNA study who met the aforementioned inclusion criteria and had complete MEG SG data were included. Thus, the final analytic sample included 32 participants with CI (Table 1).

**Table 1.**
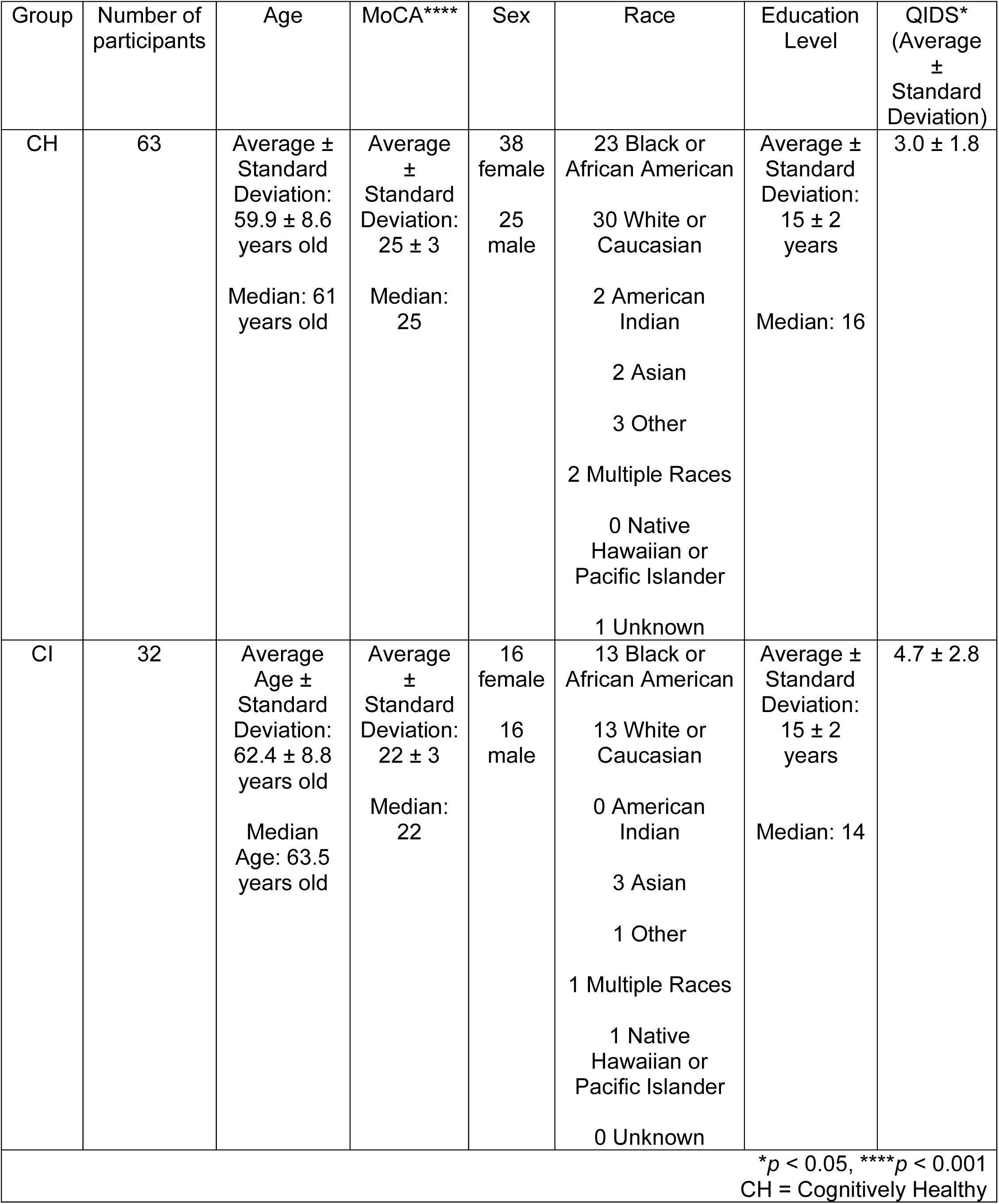
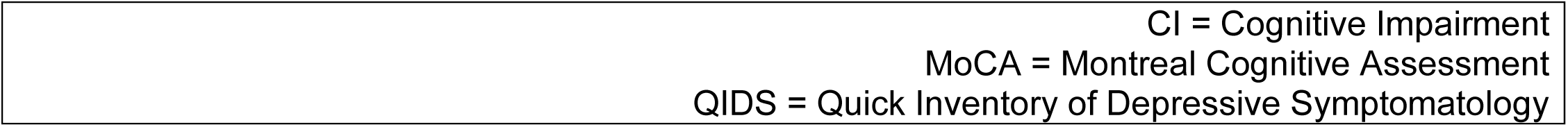
Group demographics and confounding variable (QIDS score).

| Group | Number of participants | Age | MoCA**** | Sex | Race | Education Level | QIDS*<br>(Average ± Standard Deviation) |
| --- | --- | --- | --- | --- | --- | --- | --- |
| CH | 63 | Average ± Standard Deviation:<br>59.9 ± 8.6 years old<br>Median: 61 years old | Average ± Standard Deviation:<br>25 ± 3<br>Median: 25 | 38 female<br>25 male | 23 Black or African American<br>30 White or Caucasian<br>2 American Indian<br>2 Asian<br>3 Other<br>2 Multiple Races<br>0 Native Hawaiian or Pacific Islander<br>1 Unknown | Average ± Standard Deviation:<br>15 ± 2 years<br>Median: 16 | 3.0 ± 1.8 |
| CI | 32 | Average Age ± Standard Deviation:<br>62.4 ± 8.8 years old<br>Median Age: 63.5 years old | Average ± Standard Deviation:<br>22 ± 3<br>Median: 22 | 16 female<br>16 male | 13 Black or African American<br>13 White or Caucasian<br>0 American Indian<br>3 Asian<br>1 Other<br>1 Multiple Races<br>1 Native Hawaiian or Pacific Islander<br>0 Unknown | Average ± Standard Deviation:<br>15 ± 2 years<br>Median: 14 | 4.7 ± 2.8 |

\* $p < 0.05$ , \*\*\*\* $p < 0.001$ CH = Cognitively Healthy

The non-impaired participants were further stratified into those who endorsed subjective cognitive complaints (SCC; N = 101) versus those who did not (N = 116). Specifically, the SCC participants were individuals who answered “yes” to neurocognitive pre-assessment items inquiring “Do you have any problems with your memory?” and/or “Do you have any difficulties solving problems?”. Because SCC may indicate preclinical neurodegenerative processes, participants reporting such concerns were excluded from the cognitively healthy (CH) control group (Avila-Villanueva & Fernandez-Blazquez, 2017; Rabin et al., 2017). From the CH group, additional participants were excluded due to medical comorbidities (n = 42), left-hand stimulation (n = 1), incomplete demographic data (n = 4), excessive MEG artifact (n = 3), or outlier MEG metrics (> 3 standard deviations from the mean; n = 3). Thus, the final analytic sample included 63 CH control participants (Table 1).

### Neuropsychological testing

The neurocognitive battery was administered by trained research personnel and included assessments of memory, executive function, and global cognition, including the Consortium to Establish a Registry for Alzheimer’s Disease (CERAD) word list memory, recall, and recognition (Rossetti et al., 2010), Wechsler Test of Adult Reading (WTAR) (Wechsler, 2001), Montreal Cognitive Assessment (MoCA) (Nasreddine et al., 2005), Trail Making Test (Llinas-Regla et al., 2017), Animal Fluency (Spreen, 1998), and additional behavioral measures (Haworth et al., 2016). Based on past literature (Wiesman et al., 2021), cognitive domains were constructed for (1) attention/executive function (indexed by Trail Making Test Part B [TMT-B] (Kortte et al., 2002; Llinas-Regla et al., 2017)) and (2) processing speed (derived from WAIS-IV Coding (Wechsler, 2001) and Trail Making Test Part A [TMT-A] (Llinas-Regla et al., 2017)). To account for demographic influences on cognitive performance (Gross et al., 2015; Zhang et al., 2017), raw neurocognitive scores were adjusted by regressing out the effects of sex, education, and race. Residualized scores were then z-transformed. For processing speed, z-scored residuals from WAIS-IV Coding and TMT-A were averaged to form a composite domain score. Residualized and standardized scores were used in all subsequent analyses. Age was incorporated as a covariate in the final statistical models (see *Statistical Analyses* below).

### Experimental paradigm

During MEG, participants were positioned in the 60° seated position in a nonmagnetic chair. They were instructed to close their eyes and rest their right arm on the chair tray during unilateral electrical stimulation. This stimulation was applied to the right median nerve using external cutaneous stimulators and the Digitimer DS7A constant-current stimulator system. Participants underwent 100 paired-pulse trials; thus, a total of 200 stimuli were delivered during the task. Paired-pulse trials were collected with an interstimulus interval of 500 ms and an inter-pair interval of 3950 ± 150 ms. Each pulse generated a 0.2 ms constant current square wave and was set to 10% above the motor threshold required to prompt a subtle thumb twitch. The stimulation level for each participant was recorded to assess its potential influence in regression analyses and was only reported when it demonstrated a significant covariate effect. The total run-time of the paradigm was ∼7 minutes.

### MEG data acquisition

MEG recordings were acquired using a 306-sensor MEGIN TRIUX Neo system (204 planar gradiometers, 102 magnetometers) at a sampling rate of 1 kHz and acquisition bandwidth of 0.1-330 Hz within a 3-layer magnetically shielded room. Participants were continuously monitored via real-time audio-video recording. Head position and motion were tracked using five head position indicator (HPI) coils. Prior to MEG, a 3D digitizer was used to create a map of the participant’s head including the location of fiducials and the HPI coils. During MEG acquisition, an alternating current at a unique frequency (e.g., 320 Hz) was applied to each HPI coil, generating magnetic fields that enabled continuous localization relative to the sensor array and supported offline correction of head motion. Following acquisition, MEG data were corrected for head motion and noise reduced using signal space separation with temporal extension (Medvedovsky et al., 2007; Taulu et al., 2004; Taulu & Simola, 2006).

### Structural MRI acquisition, processing, and MEG coregistration

Immediately following MEG acquisition, MRI-visible fiducial markers were placed at the nasion and preauricular points identified during the prior digitization procedure. These markers were left in place for the duration of the MRI scan to support accurate alignment between modalities. Structural images were collected on a 3T Siemens Prisma system using a 64-channel head coil. A high-resolution T1-weighted 3D Magnetization-Prepared Rapid Gradient Echo (MP-RAGE) sequence was acquired (1 mm³ isotropic voxels; repetition time = 1800 ms; echo time = 2.26 ms; field of view = 256 mm; matrix size = 256 × 256; slice thickness = 1 mm; sagittal orientation; flip angle = 8°).

Anatomical images were processed using the FreeSurfer automated recon-all pipeline (version 7.1.1). The resulting segmented volumes were then imported into Brainstorm (version 3.4) (Tadel et al., 2011) and spatially normalized to MNI space after source reconstruction (Fonov, 2009). Since MEG HPI coil locations were also known in head coordinates, these data enabled transformation of all MEG measurements into a common coordinate system. This common space was used to align each participant’s MEG data with their corresponding T1-weighted structural image prior to source-level analyses. Source reconstruction was then performed in this aligned space, after which the resulting functional images were transformed into standardized MNI space using the transformation parameters derived from the structural MRI.

To further examine potential associations with SG, structural and diffusion MRI measures were additionally evaluated, with detailed methods and diffusion MRI acquisition information presented in the Supplementary Information.

### MEG preprocessing

MEG preprocessing followed standardized procedures described in prior work (Bell et al., 2026). Data were filtered using a high-pass filter of 1 Hz, low-pass filter of 200 Hz, and a 60 Hz (and its harmonics) notch filter. Cardiac and ocular artifacts were removed using independent component analysis, and data were visually inspected to exclude residual artifact-contaminated trials (e.g., muscle). Cleaned continuous data were segmented into 4300 ms epochs (0 ms = Stimulation 1; 500 ms = Stimulation 2) with a baseline defined from −700 to −300 ms. The average number of remaining usable trials for the CH and CI groups were 87 (SD = 8) and 88 (SD = 10), respectively. Trial counts used for analysis did not differ between groups (Mann-Whitney U test, *p* > 0.05). Only data from planar gradiometers were used during analyses.

### MEG sensor-level analysis

Artifact-free epochs were transformed into the time frequency domain using Morlet wavelet analysis (range: 2-152 Hz, step: 1 Hz), averaged across trials to generate time-frequency plots of mean spectral magnitude, and baseline-normalized per sensor. The time-frequency windows used for beamforming were selected after statistical analysis of the sensor-level spectrograms across all participants, which used parametric tests against a null hypothesis that the mean is zero (thresholded at *p* < 0.05), and multiple comparisons were controlled using the Benjamini-Hochberg false discovery rate (FDR) procedure. Only time frequency windows that contained significant oscillatory responses after FDR corrections were subjected to beamforming (i.e., source level) analysis.

### MEG source-level analysis

Cortical networks were imaged via a linearly constrained minimum variance vector beamformer, which calculates source power through spatial filtering in the time-frequency domain and normalized per participant using a separately averaged baseline of equal duration and bandwidth (Hillebrand et al., 2005; Liljestrom et al., 2005; Van Veen et al., 1997). This generated 3D source maps for each time-frequency window identified in the sensor-level analysis, per participant. These images are referred to as pseudo *t*-maps that contain pseudo-t values that reflect noise-normalized power per voxel at a 4.0 x 4.0 x 4.0 mm resolution. The maps were transformed into standardized MNI space and spatially resampled to 1 mm^3^ voxels. All MEG pre-processing and imaging took place in Brainstorm (version 3.4) (Tadel et al., 2011).

The resulting voxel-wise maps of neural oscillatory response amplitude were averaged across both stimulations and all participants to generate grand-averaged beamformer images, per frequency-specific response. From each grand-averaged map the peak voxel of maximum amplitude was identified and voxel-wise time series data (i.e., virtual sensor signals) were extracted from the location, for each participant. Virtual sensor time series were derived by applying the forward model’s sensor weighting matrices to the recorded scalp-level MEG signals. The relative (baseline-normalized) time series from the peak voxel were used to compute maximum response amplitudes for the first (Stim1) and second (Stim2) stimuli, as well as the gating ratio (GR; Stim2/Stim1), per participant. To ensure numerical stability and prevent division by values near zero, frequency-specific offsets were applied prior to GR calculation. Specifically, a constant of 16 was added to theta responses and 4 to gamma responses to account for negative values, while constants of 8 and 18 were subtracted from beta and alpha responses, respectively, to adjust for positive offsets. These adjustments reflect the physiological directionality of the signals, as theta and gamma exhibit event-related synchronization (increases relative to baseline), whereas alpha and beta show event-related desynchronization (decreases relative to baseline). Using these adjusted amplitudes, GR was calculated as an index of SG. GR was selected over gating difference (GD; Stim1 − Stim2), as it accounts for inter-individual variability in response magnitude and is therefore more sensitive to amplitude-related group differences, which were previously reported in a CI cohort (Stephen et al., 2010). Absolute (non-normalized) time series were additionally extracted from the peak voxel per participant and used to quantify spontaneous activity by averaging signal amplitude within the pre-stimulus baseline period. To further characterize oscillatory dynamics, time-frequency data were smoothed at frequency- and time-specific resolutions to estimate the peak frequencies of each response (Stim 1, Stim 2) within each band: 1 Hz / 50 ms for gamma, 0.1 Hz / 250 ms for theta, 0.1 Hz / 175 ms for alpha, and 0.1 Hz / 225 ms for beta, on a per-participant basis.

### Statistical analyses

To probe for potential confounding variables between groups, demographic and clinical variables were assessed using appropriate parametric or nonparametric statistical tests for group differences. Continuous variables (e.g., age) were compared using Welch’s two-sample t-tests, which are robust to unequal variances and sample sizes, while categorical variables (e.g., sex) were evaluated using chi-square tests or Fisher’s exact tests when cell counts were less than 5. Depressive symptoms, assessed using the Quick Inventory of Depressive Symptomatology (QIDS), significantly differed between groups (*p* < 0.05) and were therefore included as a covariate in subsequent analyses to account for potential confounding effects (Rush et al., 2003) (Table 1). Age was not significantly different between groups but was also utilized as a potential confounding variable for group main effects, and a variable-of-interest for group-by-age interactions.

Our primary statistical analyses focused on evaluating group differences in MEG-derived somatosensory oscillatory metrics between CH adults and CI adults. General linear models (GLMs) were used to assess group effects on MEG metrics while adjusting for relevant demographic and clinical covariates. Age and QIDS score were included as covariates in all models, and neurocognitive performance measures were incorporated where relevant to test whether between-participant variability in neurocognitive performance masked group differences in oscillatory indices. Type III analysis of variance (ANOVA) (i.e., Type III sums of squares) was applied to all models to estimate covariate-adjusted effects of group and cognitive predictors. Interaction terms (e.g., group × age) were examined to determine whether age moderated group differences. Importantly, when interactions are included in type III ANOVAs, main effects are not interpretable; thus, main group effects were only evaluated in absence of the interaction term.

To evaluate statistical suppression effects, a regression-based approach was implemented. A reduced model (predictor + covariates) and a full model (predictor + covariates + neurocognitive variable) were compared. Indirect (ACME) and direct (ADE) effects were estimated using nonparametric bootstrapping (5,000 iterations) with bias-corrected confidence intervals using the R *mediation* package (Dustin Tingley, 2014). Statistical suppression effects were defined as cases in which inclusion of the neurocognitive variable increased the magnitude of the predictor’s direct effect and produced opposing signs between direct and indirect effects, indicating that the neurocognitive variable removed variance that obscured the predictor-outcome relationship rather than transmitting the effect (MacKinnon et al., 2000). For visualization, covariate-adjusted values (model residuals) were used to display group-level contrasts. All neurobehavioral statistical analyses were performed in R using RStudio (Posit, 2025).

## Results

Robust neural oscillatory responses to somatosensory stimulation were observed in theta (4-7 Hz; 0-250 ms post stimulation), alpha (8-13 Hz; 225-400 ms post stimulation), beta (15-25 Hz; 125-350 ms post stimulation), and gamma (30-90 Hz; 10-60 ms post stimulation) (*p* < 0.05, corrected; Figure 1a). Responses were localized to contralateral S1 in the hand-knob region (Figure 1b). Main group effects were observed in beta and gamma bands, and a group-by-age interaction was identified in the theta band, as discussed below. No significant group effects or any group-by-age interactions were observed in the alpha band.

**Figure 1.**
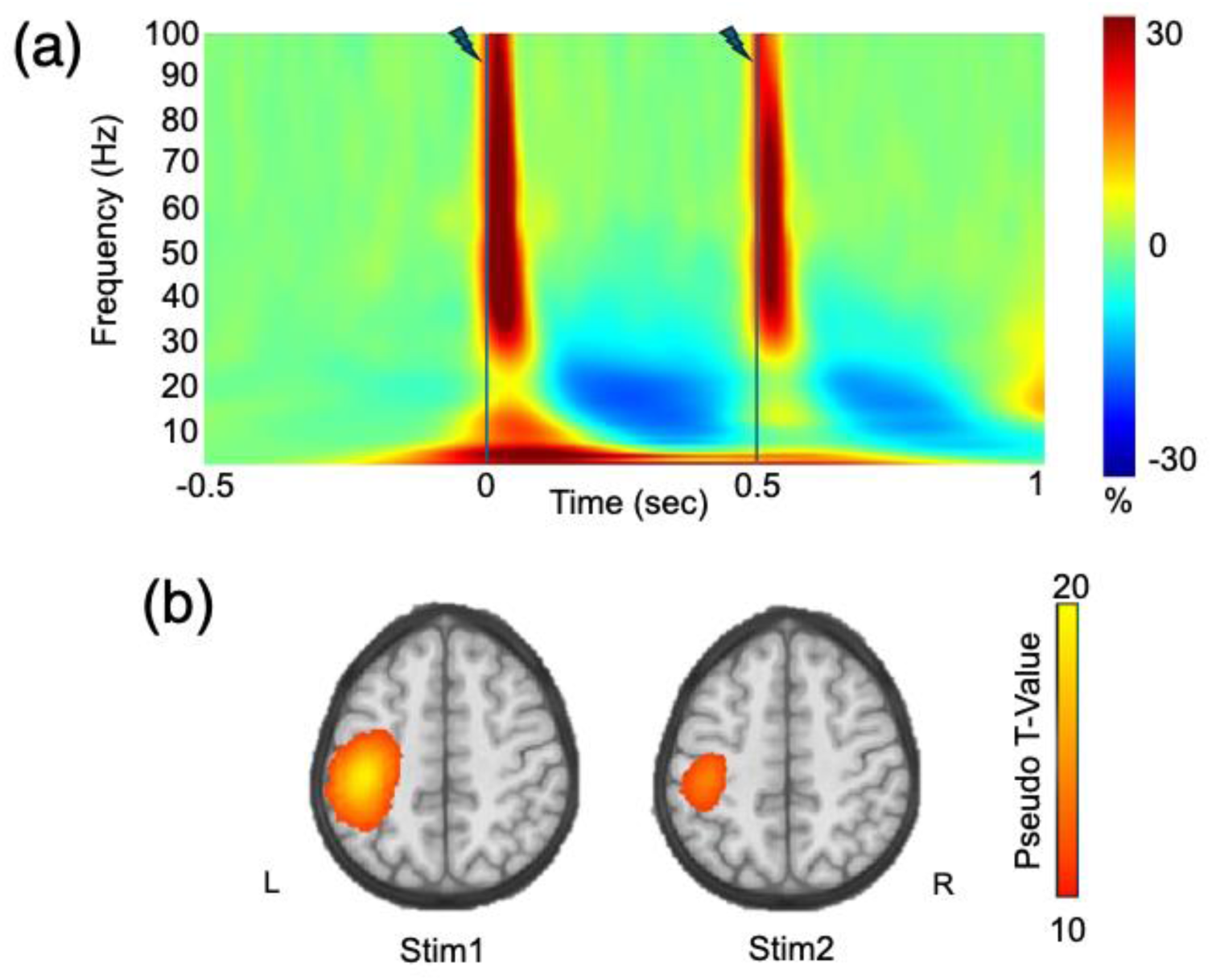
Neural Oscillatory Responses to SG Paradigm. (a) Grand-averaged time-frequency spectrogram from a MEG sensor near the contralateral sensorimotor cortex. The x-axis denotes time in seconds, beginning at -0.5 seconds and ending at 1 second. Stimulation was delivered at 0 seconds and 0.5 seconds, denoted by the line and bolt symbol. The y-axis denotes frequency (Hz), and the magnitude is shown as a percentage change unit relative to baseline period, with a color bar to the far right of the spectrogram. (b) Gamma (30-90 Hz) beamformer images (pseudo-*t* values) collapsed across both groups for Stim1 (10-60 ms) and Stim2 (510-560 ms) are shown. Robust event-related synchronizations were observed in virtually identical regions in the contralateral hand-knob area of S1 in response to each stimulation, with the response to the second stimulation being visibly attenuated, reflecting SG.

### Variability in attention/executive function masks increased gating of somatosensory gamma in CI

Given previous findings suggesting that somatosensory dysfunction is masked by cognitive impairments in Alzheimer’s spectrum patients (Wiesman et al., 2021), we evaluated whether individuals with CI display a similar trend by statistically testing main group effects on gamma SG and gamma response amplitudes with and without controlling for attention/executive function and processing speed domains in distinct models. A GLM revealed no significant group effect for gamma SG after accounting for age and depression scores (*p* = 0.145). However, when accounting for between-participant variability in attention/executive function the GLM revealed a significant group effect on gamma SG (*β* = -0.256, *t*(4, 90) = -2.076, *p* = 0.041 Figure 2b), such that individuals with CI had exaggerated gating (i.e., greater inhibition). A mediation analysis (5,000 bootstrap samples) revealed inconsistent mediation effects such that the direct effect and the mediated (i.e., indirect) effects of group on gamma SG were found to have opposite signs, making the total effect near zero and trending despite strong direct and indirect paths (Total Effect = -0.338, 95% CI [-0.721, 0.041], *p* = 0.080; Figure 2c). Specifically, the direct effect of group (ADE = -0.538, 95% CI [-0.270, -0.0010], *p* = 0.038) was significant and negative, and the indirect effect (ACME = 0.200, 95% CI [-0.048, 0.520], *p* = 0.114) was not significant and positive (Figure 2c). This finding, specifically the opposite directions for indirect and direct effects and the cancelling of the total effect, aligns with a statistical suppression effect. Thus, differences between individuals with CI and CH controls in the gating of gamma within the contralateral S1 were found to be suppressed by attention/executive function performance (i.e., Trail B time). No main group effects were found when controlling for processing speed or when evaluating gamma response amplitudes (all *p*’s > 0.05). Additionally, no main group effects were observed when evaluating theta, alpha, and beta SG and response amplitude metrics, irrespective of whether attention/executive function or processing speed performance were accounted for (all *p*’s > 0.05).

**Figure 2.**
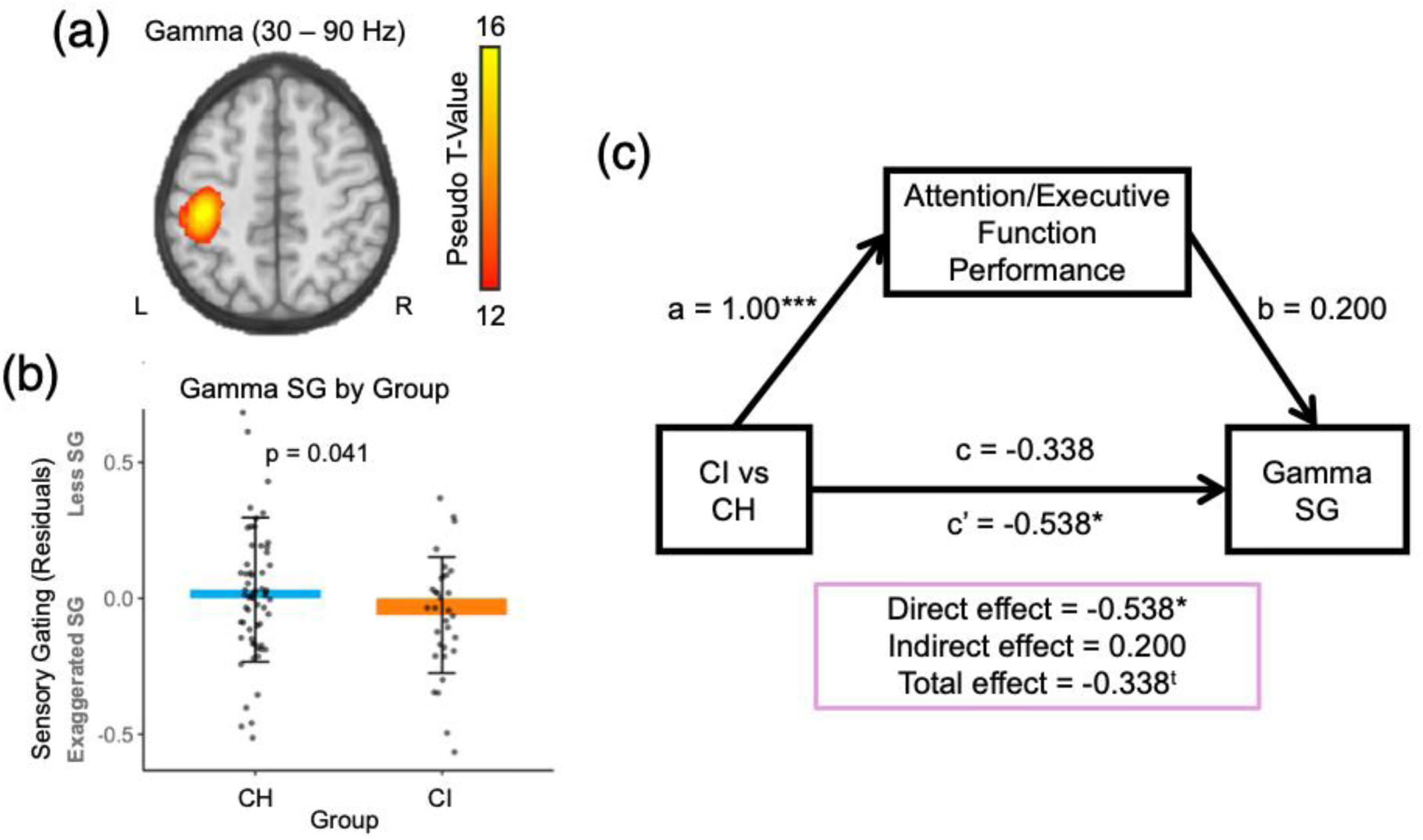
Attention/Executive Function Performance Suppresses Group Differences in Gamma SG. (a) Grand-averaged beamformer map (pseudo-*t*) for gamma (30-90 Hz) collapsed across CI and CH adults and both stimulations identified a peak voxel in the left S1. (b) When accounting for attention/executive function performance, group differences in gamma SG reach significance. Sensory gating is reported on the y-axis as residuals, above and beyond the effects of age, QIDS score, and attention/executive function performance. Bars depict adjusted means after residualizing for CH controls (blue) and individuals with CI (orange). Error bars represent ±1 standard deviation of the mean. The *p*-value reflects the group effect from a linear model including covariates and neurocognitive performance. (c) The schematic depicts a mediation framework in which group is modeled as the predictor, gamma SG as the outcome, and attention/executive function performance as the mediator. Indirect effects, direct effects, and total effects were estimated using the *mediation* package in R. Type III ANOVA were used to evaluate path coefficients. Path coefficients represent standardized regression estimates, and solid arrows denote modeled pathways between variables. CH = cognitively healthy; CI = cognitively impaired; SG = somatosensory gating; ^t^*p* < 0.10, \**p* < 0.05, \*\*\**p* < 0.005

### Somatosensory beta response peak frequency is altered in CI

Provided that frequency-specific alterations during SG have been found in other clinical populations, we explored whether the peak frequency of each oscillatory response differed by group. Using a GLM with age and nuisance variables, we observed that the peak frequency of the beta response following Stim2 (PF2) significantly differed between groups (*β* = 0.258, *t*(3, 91) = 2.500, *p* = 0.014), with PF2 elevated in CI compared to CH (Figure 3). No significant group effects were observed for beta response peak frequency following Stim1 (PF1) nor for the peak frequencies of theta, alpha, and gamma responses (all *p*’s > 0.05). Attention/executive function performance did not mediate or suppress peak frequency findings.

**Figure 3.**
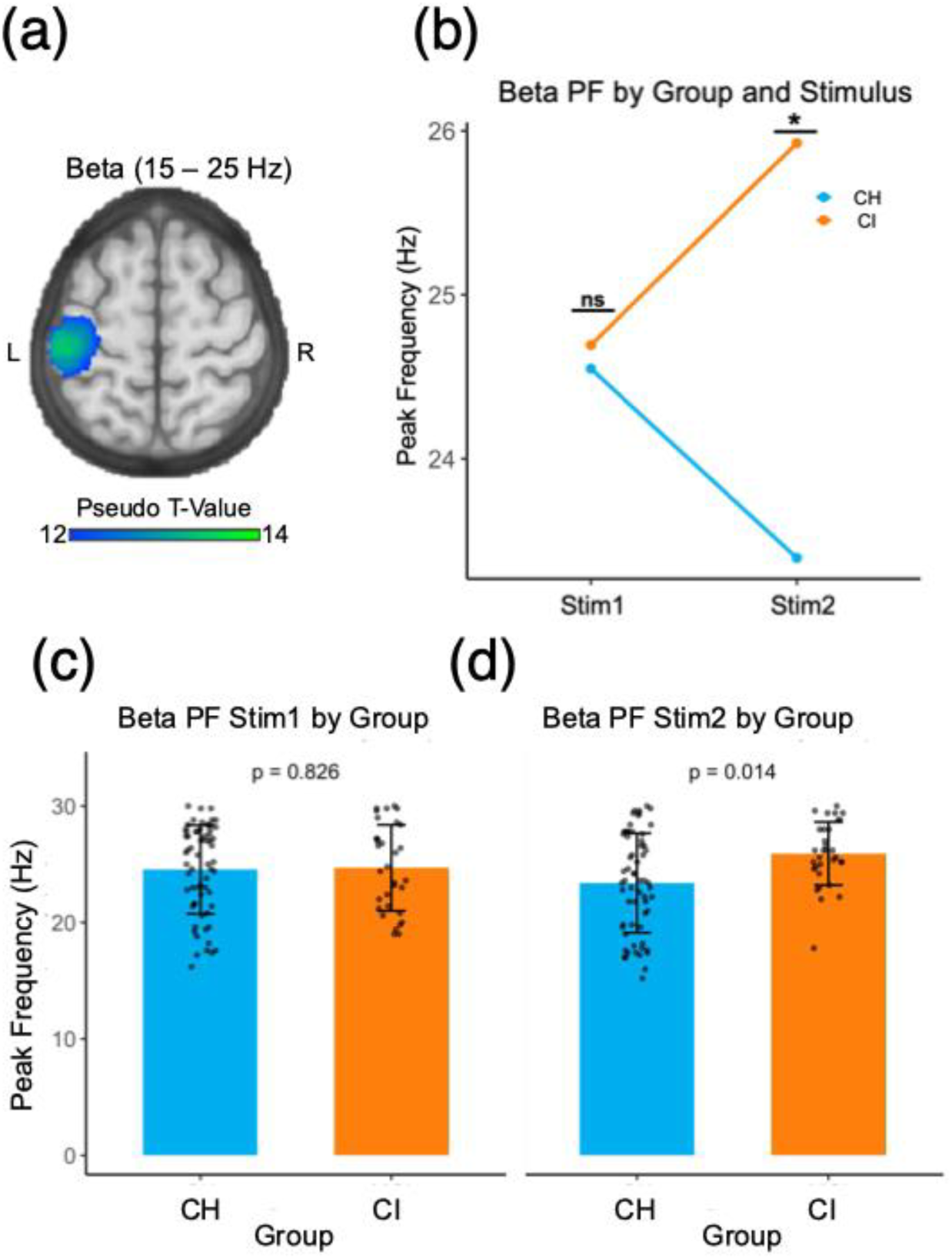
Group Differences in Beta Peak Frequency (PF) in Contralateral S1. (a) Grand-averaged beamformer map (pseudo-*t*) for beta (15-25 Hz) collapsed across CI and CH adults and both stimulations identified a peak voxel in the left S1. (b) The figure depicts group mean contralateral S1 beta response peak frequency following Stim1 and Stim2, plotted separately for individuals with CI (orange) and CH controls (blue). Values represent group-level adjusted averages (residuals) derived from separate GLMs fit for each stimulation, with age and QIDS score included as covariates. Individuals with CI had a higher beta response peak frequency following the second stimulation relative to controls. (c-d) Similar to (b), graphs represent difference in peak frequency for Stim1 and Stim2 by group. Bars depict means for CH controls (blue) and individuals with CI (orange). Error bars represent ±1 standard deviation of the mean. The *p*-value reflects the group effect from a linear model including covariates and neurocognitive performance. CH = cognitively healthy; CI = cognitively impaired; ns = not significant; PF = peak frequency; \**p* < 0.05

### Age-related changes in the gating of somatosensory theta are absent in CI

Given that prior studies have reported age-related changes in SG, we explored whether group-by-age interactions were present for SG of each oscillatory response. A GLM (with age and nuisance variables) revealed theta SG in contralateral S1 exhibited a significant group-by-age interaction (*β* = 0.217, *t*(4, 90) = 2.171, *p* = 0.033; Figure 4). Follow-up simple slopes analyses revealed that age was significantly associated with theta SG in the CH group (β = −0.048, SE = 0.014, t(90) = −3.35, *p* = .001), such that increasing age was related to exaggerated theta gating (i.e., greater inhibition). In contrast, the association between age and theta SG was not significant in the CI group (β = 0.005, SE = 0.020, t(90) = 0.25, *p* = .80). No significant group-by-age interactions were identified for gating of somatosensory alpha, beta, and gamma (all *p*’s > 0.05). Attention/executive function performance did not mediate or suppress age-related theta SG findings.

**Figure 4.**
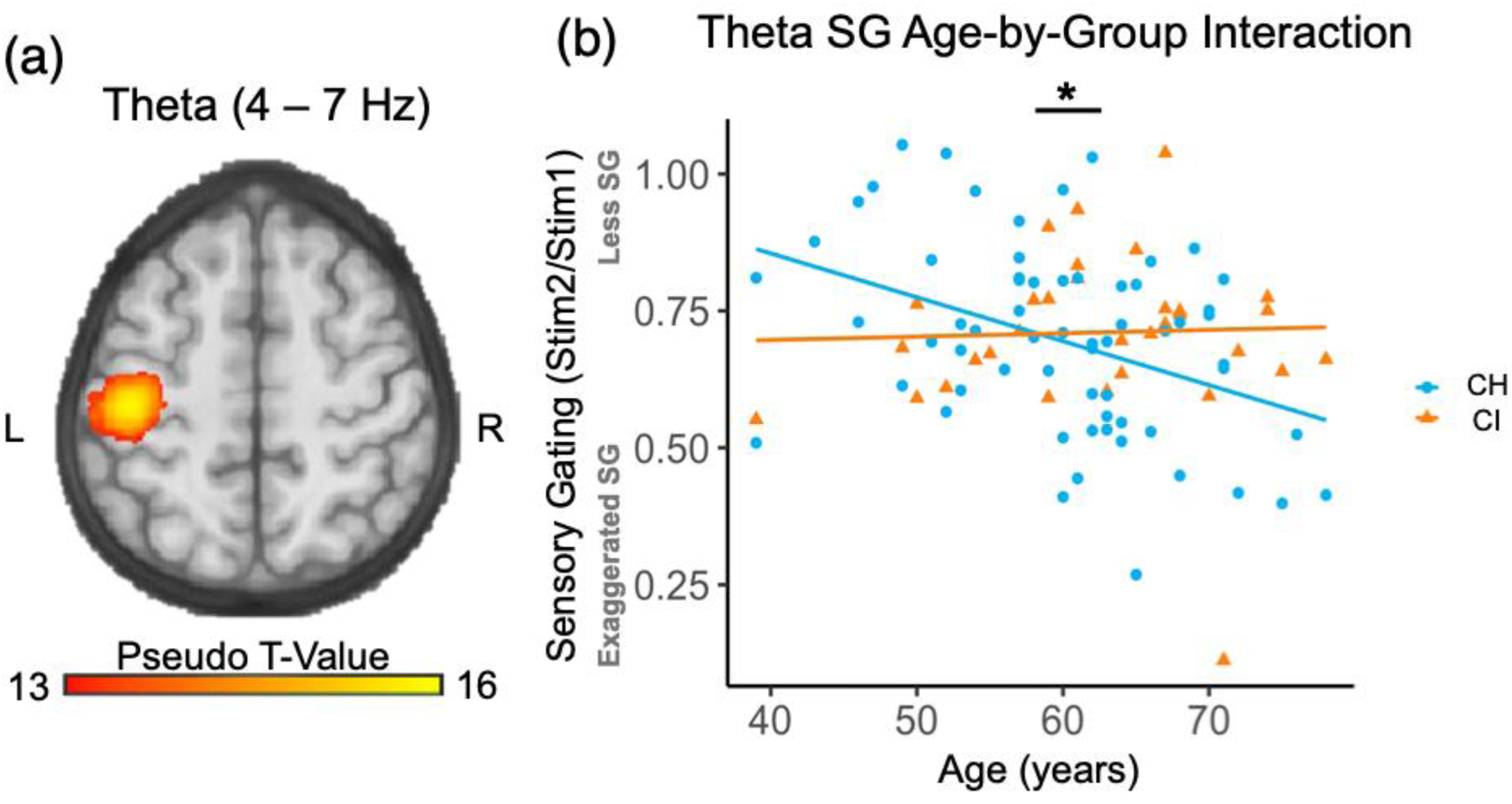
Group-by-Age Interaction in Theta SG in Contralateral S1. (a) Grand-averaged beamformer map (pseudo-*t*) for theta (4-7 Hz) collapsed across CI and CH adults and both stimulations. Following stimulation, a robust increase in theta activity was observed in a peak voxel located in the left S1. (b) The figure depicts the relationship between age and contralateral S1 theta SG plotted separately for CH (blue) and CI (orange) groups. Points represent individual participants, and lines show group-specific adjusted regression slopes derived from linear models including age and QIDS as covariates. The plotted interaction illustrates that theta SG increased with advancing age in CH adults, while age effects were not observed in adults with CI. CH = cognitively healthy; CI = cognitively impaired; SG = somatosensory gating; \**p* < 0.05

No significant group differences were observed for spontaneous theta, alpha, beta, and gamma within contralateral S1 (all *p*’s > 0.05).

Finally, structural and diffusion MRI measures were not significantly associated with the variables of interest (see Supplementary Information for results).

## Discussion

The present study addresses several limitations of prior work and expands our understanding of somatosensory dysfunction in CI. Using a neurobehavioral approach combining MEG and neurocognitive testing, we examined SG dynamics in CH adults and individuals with CI. Initial models which did not include neurocognitive performance revealed no significant group differences in gamma SG, suggesting only weak or marginal alterations in early sensory inhibitory processing. However, when attention/executive function performance were included in the model, exaggerated gamma SG emerged in the CI group compared to CH adults. Additionally, analyses across other frequency bands revealed increased beta peak frequency following Stim2 in the CI group relative to controls, as well as a group-by-age interaction in theta SG, such that SG increased with age in CH adults but not in those with CI. Together, these findings represent the first evidence of SG differences specifically in CI adults and suggest that attention/executive function performance may suppress detectable somatosensory processing deficits in earlier stages of cognitive decline.

Past studies have suggested a link between sensory processing and cognitive decline (Humes et al., 2013; Loffler et al., 2024). However, most have focused on sensory domains outside of somatosensory (i.e., visual, auditory, olfactory) (Roberts & Allen, 2016; Schubert et al., 2017). In recent years, studies on accelerated cognitive aging have suggested that cognitive impairments play a role in somatosensory dysfunction (Humes et al., 2013; Ibarra-Castaneda et al., 2025; Loffler et al., 2024; Stephen et al., 2010). Animal models and recent human studies have suggested that this sensory dysfunction is present in AD pathologies (Maatuf et al., 2016; Mueggler et al., 2003) and that these changes are suppressed by cognitive variability (Wiesman et al., 2021), while less work has been done on earlier stages of cognitive decline. Only one paper exists that examines somatosensory evoked response potentials in participants with amnestic MCI. However, that study only evaluated 4 amnestic MCI individuals, urging for larger studies (Stephen et al., 2010). Specifically, this study found that S1 event-related potentials in amnestic MCI were exaggerated with age, and additional differences in potentials between healthy controls, amnestic MCI, and Alzheimer’s disease patients were discovered after regressing out the effects on neurocognitive performance, such as attention/executive function performance (Stephen et al., 2010). However, likely due to their small sample size, neurocognitive performance was not corrected for demographic differences, which is strongly encouraged for evaluating neurocognitive performance by groups (Gross et al., 2015; Zhang et al., 2017). Additionally, that study did not investigate SG, a functional inhibitory neural phenomena, that has been found to be exaggerated in advanced stages of cognitive decline. Thus, our study design helps address several limitations of prior work and expands our understanding of somatosensory dysfunction and prodromal CI.

The most important line of evidence that influenced our study evaluated whether a mixed group of individuals with amnestic MCI or those already progressed to Alzheimer’s dementia presented with SG dysfunction, compared to CH controls (Wiesman et al., 2021). Specifically, Wiesman et al. observed exaggerated gamma SG and gamma response amplitudes in the Alzheimer’s spectrum group only when accounting for attention/executive function or processing speed. Stated differently, they found that cognitive performance (i.e. attention/executive function and processing speed) suppressed robust somatosensory processing deficits in these patients (Wiesman et al., 2021). Interestingly and mentioned above, Stephen et al. found similar response amplitude results in the time domain (i.e., evoked potentials), that were more apparent once cognitive performance was regressed out (Stephen et al., 2010). Our findings demonstrate that exaggerated gamma SG is already present at a distinct earlier stage of cognitive decline (i.e., before progression to dementia) and that cognitive variability masks this effect. Our study did not replicate the gamma response amplitude findings reported by Wiesman et al. This discrepancy may be due to our exclusion of advanced CI individuals (i.e., dementia) from our CI group, whereas Wiesman et al. a mixed group of individuals with early CI and advanced CI (i.e., amnestic MCI and Alzheimer’s dementia) (Wiesman et al., 2021). Thus, our findings suggest that somatosensory gamma response amplitude is not distinctly sensitive to early stages of cognitive decline. Further, our somatosensory findings were not suppressed by neurocognitive measures of processing speed, instead, highlighting attention/executive function performance variability as a strong correlator with masked dysfunctional somatosensory gating. Past studies corroborate this finding, suggesting that processing speed (indexed by TMT-A), does not reflect the same cognitive variability as attention/executive function performance (TMT-B) in CI (Haworth et al., 2016).

In exploratory analyses, we found that CI adults exhibited a higher peak beta frequency in response to Stim2, and not Stim1, compared to CH adults. In S1, beta peak frequency is found to correlate with levels of gamma-aminobutyric acid (GABA) in healthy adults during resting state (Baumgarten et al., 2016). Upon further testing, this could perhaps suggest a compensatory effort by the brain to engage GABA for Stim2 to preserve beta SG in individuals with CI (Spooner et al., 2018). Although peak frequency and power are inversely related, we did not see any beta response power differences in those with CI, suggesting that beta power differences are task-specific and not evident during somatosensory processing. Large multimodal neuroimaging studies are necessary to clarify these beta findings.

In additional exploratory analyses, we found a group-by-age interaction for theta SG. It appears that theta SG exaggerates with advancing age in CH individuals, but no age relationship exists in CI. Past literature using a somato-visual paired-pulse oddball paradigm suggests that theta SG exaggerates during attention to somatosensory stimulation (Wiesman & Wilson, 2020). Visual inspection of the scatterplot in Figure 4b suggests that CI adults engage theta SG at a younger age than CH adults. Furthermore, theta SG stays consistent throughout different ages in our CI group. Theta is thought to reflect somatosensory network integration and efficiency (Begus & Bonawitz, 2020; Fries, 2015; Gedankien et al., 2023; Jiang et al., 2025; Kardos et al., 2014; Kocsis et al., 2001; Wang et al., 2018; Wang, 2010; Wulff-Abramsson et al., 2025), suggesting there are alterations in how aging impacts these processes in CI. Upon further testing, this could suggest CI individuals are not engaging somatosensory efficiency or attention mechanisms appropriately and/or engaging them at younger ages. Future studies should repeat the somato-visual paired-pulse oddball paradigm in younger and older CH and CI adults to confirm whether attentional mechanisms are playing a role on the theta SG and age relationship.

Interestingly, older adults with CI are thought to have lower sensory registration (i.e., higher sensory thresholds) and lower sensory discrimination (Rhodus et al., 2022). SG is thought to reflect sensory inhibitory processing to allow for neural efficiency in processing behaviorally-relevant stimuli (Cromwell et al., 2008). This basic science perspective on SG and CI may help inform translational studies, since sensory abnormalities can be less variable than cognitive ones and further inform daily living of individuals with CI (Rhodus et al., 2022). For example, Rhodus et al. also found that sensory-based assessments and interventions gave valuable insights into behavior expression and how to improve quality-of-life for older CI adults (Rhodus et al., 2022). Specifically, one individual was found to have lower visual sensory registration, thus, urging his caretakers to use larger signs to remind him to drink water and find his keys (Rhodus et al., 2022). Further studies should evaluate whether somatosensory processing (i.e. texture of clothing, heat discrimination, social tactile contact) is altered in older adults with CI and whether interventions considering somatosensory impairments can improve their daily quality of life.

Our study has several limitations that should be considered when interpreting these findings. First, we were unable to stratify individuals with CI based on Alzheimer’s biomarker status or clinical diagnoses of amnestic MCI. As a result, the present sample likely includes a heterogeneous group of individuals with varying underlying etiologies of CI. Future studies incorporating biomarker confirmation (e.g., amyloid or tau markers) will be important for determining whether the observed SG differences are specifically associated with MCI due to Alzheimer’s pathology. Second, a key difference between previous studies and the current investigation is the defining criteria for CI. Both Stephen et al. and Wiesman et al. focused on individuals with amnestic MCI, which primarily reflects memory decline (Stephen et al., 2010; Wiesman et al., 2021). In contrast, our CI group included individuals with more diverse cognitive impairments rather than restricting the sample to amnestic presentations. While this approach increases generalizability, it may also introduce additional heterogeneity in the cognitive and neural profiles represented in the sample. Finally, because the sample size was modest and depressive symptoms (QIDS scores) were included as a covariate, inferences regarding group-by-age interactions should be interpreted cautiously. Future studies with larger samples will be necessary to provide greater statistical power to more robustly evaluate these effects.

## Conclusion

In conclusion, gamma SG exaggeration, increased beta response PF following Stim2, and lack of age and theta SG correlations suggests early, aberrant somatosensory processing and inhibition in the CI group. Importantly, CI reflects accelerated aging and, in some cases, can progress to dementia or Alzheimer’s disease (Anand S, 2025; Collie & Maruff, 2000). Somatosensory dysfunction in Alzheimer’s disease is thought to be spared until later stages of disease progression (Abbruzzese et al., 1984; Ewers et al., 2011; Frisoni et al., 2010; Uylings & de Brabander, 2002). Our findings suggest that somatosensory processing deficits are present at earlier, less severe stages of CI in adults and at times reflect a different aging trajectory than healthy controls. Follow-up studies are needed to confirm subcategories of MCI diagnoses and see if SG metrics can differentiate amnestic or non-amnestic MCI. Finally, and most importantly, future studies should investigate somatosensory processing, along with other sensory modalities, when drawing conclusions about sensory impairments and cognitive dysfunction. Our results coupled with those within the larger field demonstrate bi-directional impacts of somatosensory processing and cognition in healthy and disease states, and these relationships should be further studied to uncover translational potential.

## Data statement

Requests for access to DHMS data may be submitted to the Dallas Hearts Study (DHS) Publications Committee according to established study procedures.

## Author Contributions

ALP and EMD contributed to the development, design, and validation of the MEG methodology. FFY contributed to the development and design of the MRI methodology. LHL, HR, and CMC contributed to the development and design of the neuropsychological methodology. MV completed the formal analyses, visualization, investigation, and wrote the original draft of the manuscript. MV, LG, and SW contributed to MRI data preprocessing, and ALP, EMD, MV, NMB, LG, and SW assisted in data collection. YX, FFY, AMS, JAM, and ALP contributed to the supervision of the data analyses. YX, NMB, TP, LG, SW, FFY, LHL, HR, CMC, AMS, EMD, JAM, and ALP provided edits to the manuscript.

## Acknowledgements

We would like to acknowledge Adriana Ohm in her assistance with DHMS piloting and data collection. The DHMS at the UTSW Medical Center was funded through the Harry S. Moss Heart Trust. The funders had no role in study design, data collection and analysis, decision to publish, or preparation of this manuscript.

## Notes

### Competing Interest Statement

The authors have declared no competing interest.

