## Supplemental materials for "Somatosensory gating dysfunction is masked by cognitive variability in cognitively impaired individuals"

### Supplementary Materials

#### Methodology for surface-based morphometry analyses

Past structural MRI studies of cognitive impairment (CI) have demonstrated cortical thinning with widespread neocortical involvement observed in some samples, including sensorimotor areas (Chen et al., 2024). However, such effects are typically diffuse and variable (Singh et al., 2006; Sun et al., 2019), and the primary somatosensory cortex (S1) is not consistently identified as a selective or early site of structural degeneration in CI (Chen et al., 2024; Nickl-Jockschat et al., 2012; Singh et al., 2006; Sun et al., 2019). More consistently, advanced stages of Alzheimer's disease present with structural alterations in S1 (Frisoni et al., 2010; Singh et al., 2006; Uylings & de Brabander, 2002).

Cortical thickness and gray matter (GM) volume were estimated using the automated recon-all pipeline in FreeSurfer (version 7.1.1) (Fischl, 2012). This processing stream includes intensity normalization, skull stripping, white-gray matter segmentation, and surface reconstruction from each participant's high-resolution T1-weighted anatomical MRI. Cortical thickness was computed as the distance between the white matter and pial surfaces at each vertex, while GM volume was derived from the parcellated anatomical segmentation produced during the reconstruction steps. Vertex-wise thickness and volume estimates were parcellated into anatomical regions according to the Desikan–Killiany–Tourville atlas to generate region-of-interest (ROI) values for the left-hemisphere postcentral gyrus. Because GM volume is strongly influenced by head size, all models that included volume incorporated the total intracranial volume as a covariate. ROI-extracted postcentral gyrus cortical thickness and GM volume values were entered into linear regression models testing associations with group. If group effects were present, volumetrics were also tested against magnetoencephalography (MEG)-derived somatosensory metrics.

In contrast to the robust group differences observed in MEG-derived somatosensory gating (SG) metrics, no significant group differences were detected in left postcentral gyrus cortical thickness or volume (all *p*'s > 0.05). This dissociation suggests that the observed functional alterations are unlikely to be driven by gross structural differences in S1. Volumetric measures provide relatively coarse indices of cortical architecture and may be less sensitive to microcircuit-level alterations, such as changes in excitation-inhibition balance or neuronal synchrony, that are more directly reflected in oscillatory dynamics measured with MEG. Thus, these findings suggest that group differences in somatosensory processing primarily reflect functional rather than structural alterations within primary sensory cortex. Further, the lack of structural alterations in the CI group in this study might suggest that functional alterations precede structural changes in S1. However, the CI sample size was modest, and more minute structural changes might require a larger sample size for appropriate power.

#### Methodology for diffusion analyses utilizing fractional anisotropy

Past studies have highlighted changes in fractional anisotropy (FA) as a potential variable of interest in the diagnostic processes detecting CI and dementia (Hall et al., 2021). Herein, diffusion-weighted images were acquired using a 102-direction multi-shell sequence with b-values of 0, 500, 1000, 2000, and 3000 s/mm$^2$, along with an additional reversed phase-encoding b0 image to facilitate correction of magnetic susceptibility-induced distortions. Diffusion data were preprocessed using the FMRIB Software Library (FSL; version 6.1, https://fsl.fmrib.ox.ac.uk), including topup, eddy current and motion correction (Jenkinson et al., 2012). Following preprocessing, diffusion tensors were fitted at each voxel using FSL's dtifit to generate FA maps.

Voxel-wise analyses of white matter microstructure were conducted using Tract-Based Spatial Statistics (TBSS) within FSL. All subjects' FA images were nonlinearly aligned to the FMRIB58_FA standard-space template and resampled to 1 mm isotropic resolution . A mean FA image was generated and thinned to create a white matter skeleton representing the centers of common tracts (thresholded at FA > 0.2), onto which each participant's FA data were projected. Voxel-wise statistical analyses were performed using permutation-based nonparametric testing (randomise; 5,000 permutations). The main effect of group was modeled with age and Quick Inventory of Depressive Symptomatology (QIDS) scores included as covariates of no interest (Rush et al., 2003). A separate model was used to evaluate the group-by-age interaction. Threshold-free cluster enhancement was applied, and results were corrected for multiple comparisons using family-wise error correction at $p < 0.05$.

To further examine relationships with functional measures, mean FA values were extracted from significant clusters and correlated with MEG-derived metrics. These exploratory analyses did not yield significant associations (all *p*'s > 0.05). These findings suggest that functional changes in CI were not correlated with changes in white matter integrity. However, due to our small sample size, multimodal imaging analyses should be repeated in a larger sample to confirm null findings.
